# Multi-tissue epigenetic analysis of the osteoarthritis susceptibility locus mapping to the plectin gene *PLEC*

**DOI:** 10.1101/2020.01.28.917401

**Authors:** A.K. Sorial, I.M.J Hofer, M. Tselepi, K. Cheung, E. Parker, D.J. Deehan, S.J. Rice, J. Loughlin

**Affiliations:** Newcastle University, Biosciences Institute, Newcastle upon Tyne, UK; Freeman Hospital, Newcastle Upon Tyne, UK

**Keywords:** osteoarthritis, genetic risk, epigenetics, gene expression, PLEC, CRISPR/Cas9, cytoskeletal

## Abstract

**Objective:** Osteoarthritis (OA) associated single nucleotide polymorphism (SNP) rs11780978 correlates with differential expression of *PLEC*, and methylation quantitative trait loci (mQTLs) at *PLEC* CpGs in cartilage. This implies that methylation links chondrocyte genotype and phenotype, thus driving the functional effect. *PLEC* encodes plectin, a cytoskeletal protein that enables tissues to respond to mechanical forces. We sought to assess whether *PLEC* functional effects were cartilage specific.

**Method:** Cartilage, fat pad, synovium and peripheral blood were collected from patients undergoing arthroplasty. *PLEC* CpGs were analysed for mQTLs and allelic expression imbalance (AEI) was performed. We focussed on previously reported mQTL clusters neighbouring cg19405177 and cg14598846. Plectin was knocked down in a mesenchymal stem cell (MSC) line using CRISPR/Cas9 and cells phenotyped by RNA-sequencing.

**Results:** Novel mQTLs were discovered in fat pad, synovium and peripheral blood at both clusters. The genotype-methylation effect of rs11780978 was stronger in cg14598846 than in cg19405177 and stronger in joint tissues than in peripheral blood. We observed AEI in synovium in the same direction as for cartilage. Knocking-down plectin impacted on pathways reported to have a role in OA, including Wnt signalling, glycosaminoglycan biosynthesis and immune regulation.

**Conclusions:** Synovium is also a target of the rs11780978 OA association functionally operating on *PLEC*. In fat pad, mQTLs were identified but these did not correlate with *PLEC* expression, suggesting the functional effect is not joint-wide. Our study highlights interplay between genetic risk, DNA methylation and gene expression in OA, and reveals clear differences between tissues from the same diseased joint.

## Introduction

Osteoarthritis (OA) is an extremely common musculoskeletal disorder that is characterised by focal loss of articular cartilage leading to chronic pain, disability, co-morbidities including cardiovascular disease, and premature death^1–3^. OA is polygenic and in recent years over 90 genome-wide significant loci have been mapped in Europeans using high-density single nucleotide polymorphism (SNP) arrays^4–10^. An overwhelming majority of the associated polymorphisms reside in non-protein-coding regions of the genome and, as such, OA genetic susceptibility is presumed to act principally via changes to gene expression. The availability of excised tissue following arthroplasty of an OA joint offers the opportunity to experimentally test this in disease-relevant cells, including cartilage chondrocytes, the single cell type in this tissue^11^. As such, many OA risk alleles have been shown to correlate with alteration in the expression of nearby genes in chondrocytes^12^. These effects are known as expression quantitative trait loci (eQTLs). DNA methylation changes at CG dinucleotides (CpGs) also regulate gene expression and OA-associated SNPs have been shown to correlate with methylation levels in cartilage DNA in *cis*^13^. These effects are known as methylation QTLs (mQTLs). Furthermore, several OA risk loci correlate with both methylation and expression, with alteration of the former being shown experimentally to modulate the latter^14^. These effects are known as methylation and expression QTLs (meQTLs). Overall, these studies indicate that DNA methylation links chondrocyte genotype and phenotype, with this epigenetic mechanism underpinning and driving a large proportion of the functional impact of OA genetic risk. Furthermore, in most instances, OA mQTLs localize to predicted enhancers^13^, placing OA in the “enhanceropathy” category of common diseases^15^.

We previously reported that an OA association marked by SNP rs11780978 (G>A) correlated with methylation of nine CpGs measured using the Illumina Infinium HumanMethylation450 genome-wide CpG array^16^. The SNP and CpGs are all located within the *PLEC* gene on chromosome 8q24.3. We subsequently focused on two of the most significant CpGs from amongst the nine, cg19405177 and cg14598846, and by using pyrosequencing, we replicated the correlation between rs11780978 genotype and CpG methylation in an independent panel of DNAs. The two pyrosequencing assays used to replicate cg19405177 and cg14598846 captured an additional seven CpGs flanking cg19405177 and an additional three CpGs adjacent to cg14598846. The methylation of these additional CpGs also correlated with rs11780978 genotype. Furthermore, using allelic expression imbalance (AEI) analysis (a complementary approach to standard eQTL analysis^17^) we demonstrated that the OA risk-conferring A allele of rs11780978 correlated with reduced *PLEC* expression, and that expression of the gene also correlated with methylation at CpGs in the cg19405177 and cg14598846 clusters^16^. These mQTL, eQTL and meQTL studies were all performed on cartilage DNA and RNA, and highlighted *PLEC* as a target of the OA susceptibility mapping to chromosome 8q24.3.

*PLEC* encodes plectin, a large cytoskeletal protein that regulates signalling from the extracellular environment to the cell nucleus^18–19^. Plectin enables cells to respond to external mechanical stimuli and forces, such as those experienced by the chondrocyte during joint movement^20^. Although cartilage loss is central to the OA disease process, there are pathological changes to other tissues of the articulating joint that occur concurrent to and following the loss of cartilage^21^. *PLEC* is expressed widely and in this report, we set out to test whether the mQTL, eQTL and meQTL effects that we had discovered were cartilage specific or whether they were active in other joint or non-joint tissues. We also modelled the transcriptional effect that the OA risk-conferring allele of rs11780978 has on *PLEC* by knocking-down plectin in a mesenchymal stem cell line using CRISPR/Cas9 and assessing the impact on global gene expression.

## Materials and Methods

### OA Patients

Tissue samples were collected from patients undergoing knee or hip arthroplasty for primary OA at the Newcastle upon Tyne Foundation Trust hospitals. Ethical approval was granted by the Health Research Authority of the National Health Service (research ethics committee reference 14/NE/1212) and each donor provided verbal and written informed consent. We collected macroscopically intact cartilage (36 samples, distal from the lesion and avoiding areas of fibrillated tissue), infrapatellar fat pad (68 samples) and synovium (81 samples). We also collected whole peripheral blood samples from 55 of the patients just prior to their surgery, using EDTA vacutainers for DNA extraction and Tempus™ tubes for RNA extraction (ThermoFisher Scientific). In total, we analysed 240 samples from 202 OA patients, with some patients donating more than one tissue type for analysis. Further patient details can be found in Supplementary Table 1.

### Nucleic acid extraction from tissue samples

Joint tissue samples were stored frozen and ground to a powder using a mixermill (Retsch Limited) under liquid nitrogen. For cartilage, RNA was extracted using TRIzol (Life Technologies) and DNA was extracted using the E.Z.N.A. Tissue DNA isolation kit (Omega Biotek, VWR). For fat pad and synovium, DNA and RNA were extracted using the E.Z.N.A. DNA/RNA isolation kit (Omega Biotek, VWR). For blood, DNA was extracted using the QIAamp DNA blood mini kit (Qiagen) and RNA was extracted using the Tempus™ Spin RNA isolation kit (ThermoFisher Scientific).

### Quantitative gene expression

cDNA synthesis and quantitative PCR (qPCR) were performed as described previously^22^. Predesigned TaqMan assays (Integrated DNA Technologies) were used to quantify expression of the housekeeping genes *HPRT1*, *18S* and *GAPDH*. For *PLEC*, expression primers were designed using the Roche probe library system and are located in exons 19 and 20, which are present in all isoforms of the gene (primers 5’-CTGCGTAGGAAATACAGTT-3’ and 5’-CAGCTGTTCCTTCTCGTCCT-3’). The relative expression of *PLEC* was calculated by the 2^−ΔCt^ method, where ΔCt is the mean Ct value of the three housekeeping genes subtracted from the Ct value of *PLEC*.

### mQTL and eQTL analysis

Genotyping of association SNP rs11780978 and *PLEC* transcript SNP rs11783799 (A>G), DNA methylation analysis at the cg19405177 and cg14598846 CpG clusters, and *PLEC* AEI analysis using rs11783799 were all performed as described previously using the same protocols and assays^16^. In the methylation analysis, PCRs were performed in duplicate on each sample and the mean calculated, with samples being excluded from the analysis if the methylation between the PCR replicates differed by >5%. In the AEI analysis, PCRs were performed in triplicate on each tissue cDNA and DNA sample, with samples excluded from the analysis if the values between the PCR replicates differed by >5%.

### Generation of a mesenchymal stem cell line constitutively expressing Cas9 (MSC-Cas9)

We purchased the immortalised human mesenchymal stem cell (MSC) line SCRC-4000 (ATCC) and transduced it with the lentiCRISPR v2 plasmid 52961, which contains a Cas9 insert and confers puromycin resistance (Addgene). The cells were expanded in complete growth medium consisting of serum free media (PCS-500-030, ATCC) supplemented with Mesenchymal Stem Cell Growth Kit containing L-Alanyl-L-Glutamine (PCS-500-040, ATCC), G418 disulphate salt solution (G8168, Sigma Aldrich) and penicillin-streptomycin (30-2300, ATCC). Selection of Cas9 expressing cells was achieved by addition of 0.35ug/μl of puromycin to the complete growth media. Constitutive expression of Cas9 was Confirmed by immunoblot with a Cas9 antibody (7A9-3A3, Cell Signalling Technology).

### CRISPR/Cas9

CHOPCHOP (https://chopchop.cbu.uib.no/) was used to design guide RNA (gRNA) sequences that would generate a 26bp frame-shift deletion of exon 3 of *PLEC* (primers 5’-CTGTATGAAGACCTCCGCGA-3’ and 5’-CCCCGAGAGGACCTCCAGCA-3’). This exon is present in all isoforms of the gene. The gRNAs were then purchased as synthetic Alt-R® CRISPR-Cas9 crRNAs alongside Alt-R® CRISPR-Cas9 tracrRNA (Integrated DNA Technologies). MSC-Cas9 cells were plated at sub-confluence (45,000 cells/well) in 6-well plates (VWR) and allowed to adhere for 24 hours. For deletions (4 replicates), paired crRNAs were annealed to tracrRNA in a 1:1:2 ratio at 95°C for 5 minutes, whilst for controls (4 replicates), tracrRNA alone was used. Deletion and control mixes were separately combined with Dharmafect1 (Dharmacon) in serum free media (PCS-500-030, ATCC) to produce a transfection mix. 200ul of media/well was aspirated and replaced with transfection mix for 24 hours, after which the media was changed to complete growth medium and the cells expanded in T182.5 flasks (VWR) in monolayer and until confluent (between 10-14 days), with two changes of media per week. Cells were detached by trypsinization, washed in phosphate-buffered saline (PBS), spun and the cell pellets frozen. DNA, RNA and protein were simultaneously extracted from pellets using the NucleoSpin TriPrep Kit (Macherey-Nagel, supplied by ThermoFisher). Deletion of the target region was confirmed by end-point of DNA using primers flanking the guides (primers 5’-AGGGGTTGAGCTGGATTCC-3’ and 5’-CCAGCACCCACTCTGTAGAT-3’) and by Sanger sequencing.

### Immunoblotting

Up to 10μg of total protein was resolved on NuPAGE™ 3-8% Tris-Acetate Protein Gels (Invitrogen) in 1x NuPAGE™ Tris-Acetate SDS Running Buffer (Invitrogen). Protein was transferred through a semi-dry transfer at 200mA/gel for 3 hours. Blots were probed with anti-plectin guinea pig polyclonal serum (GP21, Progen)/goat anti-guinea pig (AB6908, Abcam) and anti-GAPDH (MAB374, Sigma Aldrich)/goat anti-mouse (P0447, Dako) as a loading control.

### RNA-sequencing

For each sample, the TruSeq Stranded mRNA library prep kit (Illumina) was used to prepare a sequencing library from 1.5μg of total RNA. All samples were then sequenced to a depth of ~52-71 million paired end reads on an Illumina NextSeq platform. Quality control of raw sequencing reads was performed using FastQC (v0.11.7). All samples passed QC. Reads were mapped to the hg38 (ENSEMBL version 91) reference genome using Salmon (v0.14.1) in quasi-mapping mode on default settings^23^. All samples except one achieved alignment rates of ~88%. Counts were summarised to gene level and imported into R using tximport^24^. Principle component analysis (PCA) revealed the sample with the low alignment rate to be an outlier and this was removed from further analysis, leaving four *PLEC* deletion and three controls. Unwanted variation in the dataset was estimated using surrogate variable analysis (svaseq^25^). The number of surrogate variables was estimated at two and these were included into the model for differential expression analysis. Differential expression analysis between deletion and control samples was performed using the Wald test implemented in DESeq2^26^. The false discovery rate (FDR) was controlled using the Benjamini-Hochberg procedure. Genes with an FDR *P*<0.05 were considered significant. Gene set enrichment analysis (GSEA^27^) was used to discover overrepresented pathways in the set of up- and down-regulated genes. The full gene list was ranked by signed log10(*P* value); positive sign if up-regulated and negative if down-regulated. This pre-ranked list was used as input into the fgsea package in Bioconductor to query enrichment of Reactome pathways^28^. Pathways with an FDR *P*<0.25 were considered significant, as recommended by the developers of GSEA^27^. To visualise splice junctions, reads were aligned to hg38 reference genome using STAR (v2.4.2a) short read aligner using default settings^29^ and viewed in Integrative Genomics Viewer (IGV^30^).

### Statistical analyses

When correlating association between rs11780978 genotype and methylation of CpGs, *P* values were calculated using the Kruskal-Wallis test. For comparison of global methylation between tissues, Kruskal-Wallis with Dunn’s multiple comparisons was used. For quantitative *PLEC* expression analyses, *P* values were calculated using a Mann-Whitney 2-tailed exact test. For *PLEC* AEI, *P* values were calculated using Wilcoxon’s matched pairs signed rank test. For heat maps, the correlation between all three genotypes (GG, GA and AA) and methylation was determined using adjusted r^2^ values calculated using a standard least squares linear regression model and expressed as a percentage.

## Results

### *PLEC* cartilage mQTLs are present in other tissues

We previously discovered *PLEC* mQTLs in cartilage DNA^16^. For cg19405177 and its seven flanking CpGs (located in a 124bp interval), the OA risk-conferring A allele of rs11780978 correlated with increased methylation, whereas for cg14598846 and its three adjacent CpGs (located in a 21bp interval), the A allele correlated with decreased methylation. Supplementary Figure 1 highlights the relative positions of rs11780978 and the CpGs in *PLEC*. In this report, we first assessed if these mQTLs were also active in DNA from fat pad, synovium and blood. We also included a new cohort of cartilage DNAs. Figure 1 provides the data for cg19405177 and cg14598846, whilst the data for all 12 CpGs across the four tissues can be found in Supplementary Figures 2 (cartilage), 3 (fat pad), 4 (synovium) and 5 (blood). The mQTLs were replicated in the new batch of cartilage DNAs and in the same directions as before. The mQTLs were also present in the other tissues and also in the same direction.

**Figure 1.**
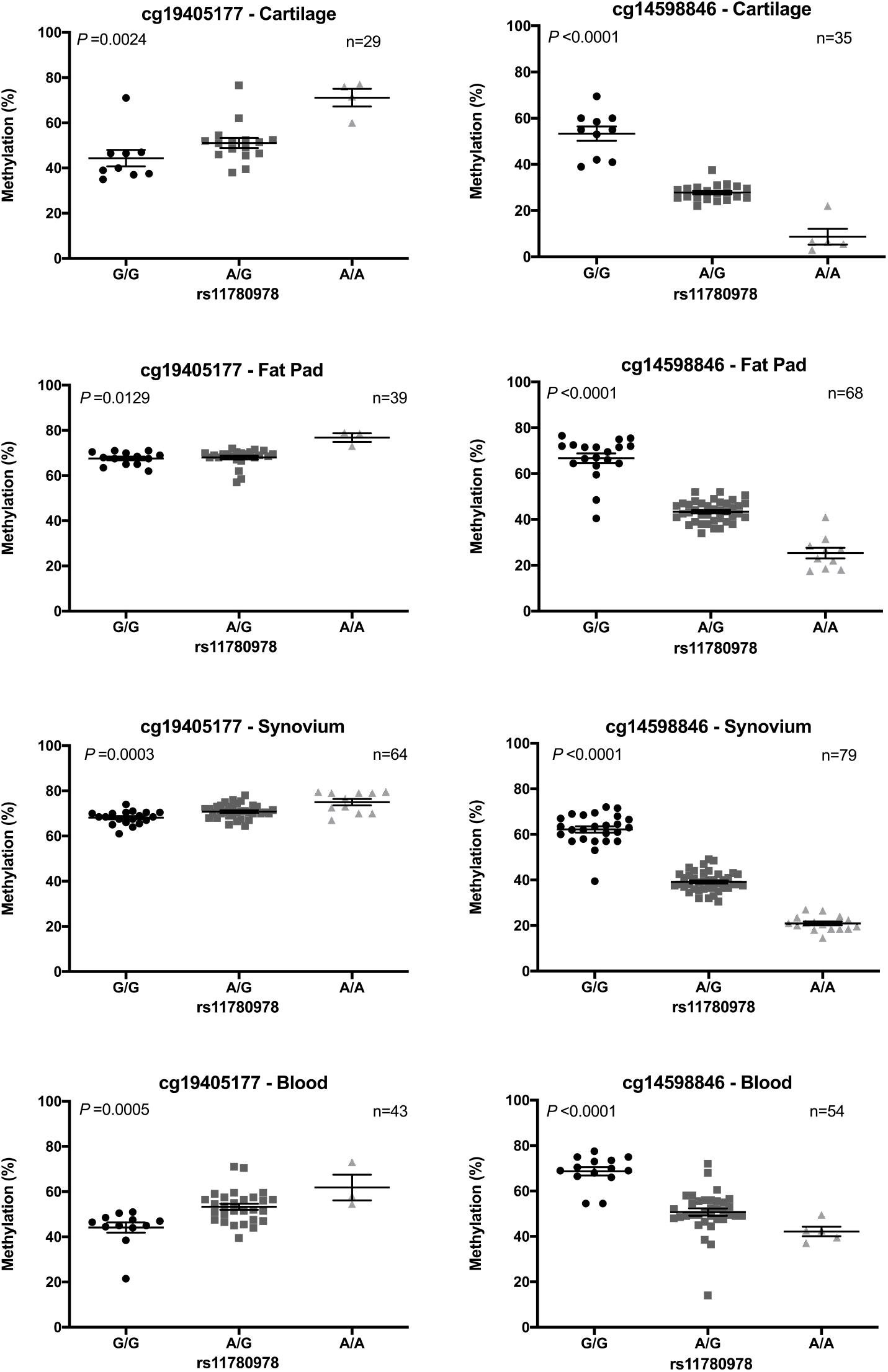
Association between rs11780978 genotype and methylation at CpGs cg19405177 and cg14598846 in cartilage, fat pad, synovium and blood DNA. *P* values were calculated using the Kruskal-Wallis test. Horizontal lines and error bars show the mean ± SEM. n = the number of patients providing data per CpG site for each tissue.

When we plotted the methylation data by tissue type but independent of genotype, there were significant differences in methylation levels (Supplementary Figure 6). For example, cg19405177 demonstrated higher methylation values in fat pad and synovium compared to cartilage and blood, whilst cg14598846 demonstrated higher methylation values in blood compared to joint tissues. To emphasize this variability in CpG methylation in tissues from the same patient, we plotted the cg19405177 and cg14598846 data for those patients in whom we had studied two or more tissues. There were 18 such patients with data generated at cg19405177 and 20 patients with data generated at cg14598846 (Figure 2). The variability in methylation between the tissues of an individual is particularly clear for patient 6550 at cg19405177, and patient 6599 at cg14598846, with a methylation range of >30% across the four tissues tested for each CpG.

**Figure 2.**
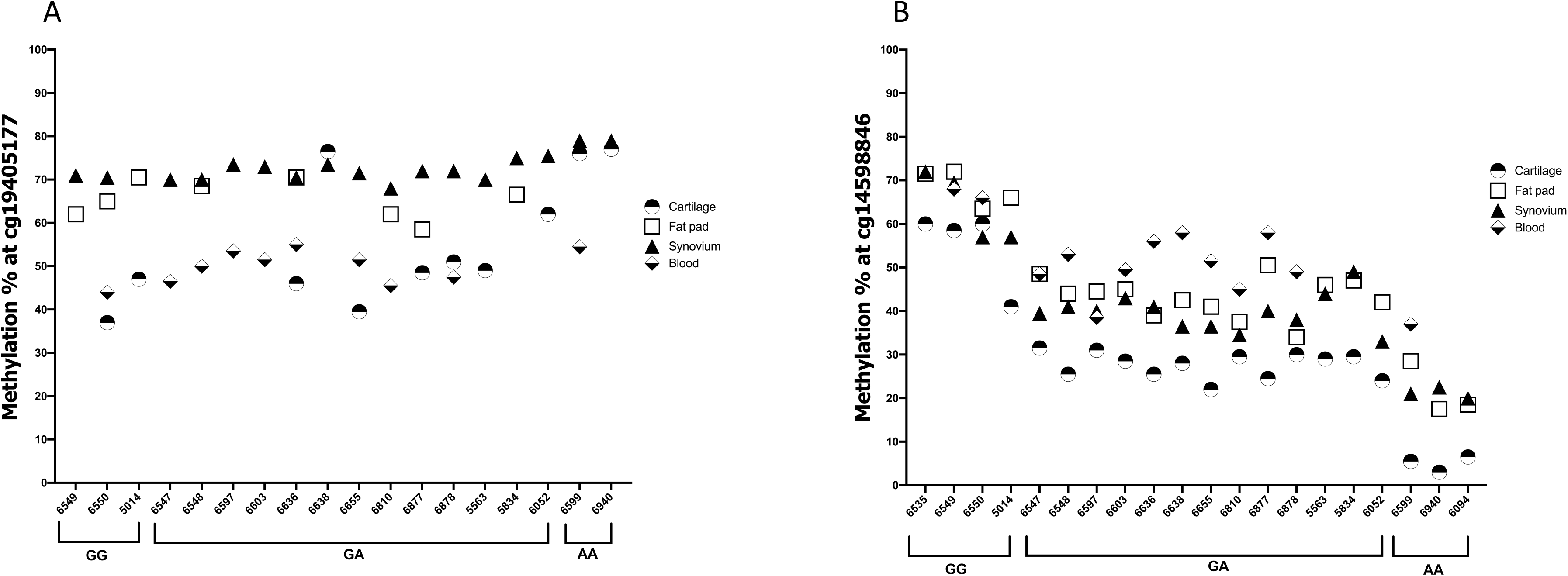
Differences in methylation within tissues of the same patient. Methylation at CpGs cg19405177 (**A**) and cg14598846 (**B**) were plotted for those individuals in whom methylation data was generated for two or more tissues. Numbers on the x-axis refer to the anonymised identification number assigned to patients at recruitment. Data is grouped by genotype at rs11780978.

We next constructed heat maps of the percentage effect of rs11780978 genotype on methylation at each CpG in each tissue. For cartilage, we included data from the replication study of our previous report^16^ to maximise power. For the cg19405177 CpGs, the average genotypic effect for all eight CpGs combined was 31.3% in cartilage, 17.4% in fat pad, 21.2 % in synovium and 9.3% in blood (Figure 3A). For the cg14598846 CpGs, the average genotypic effect for all four CpGs combined were comparable across the joint tissues, with values of 85.0% in cartilage, 81.4% in fat pad and 87.5% in synovium (Figure 3B). Although the genotypic effect in blood was lower, it was still high at 43.5%. At the individual CpG level, there were some striking differences between tissues. For example, cg19405177 had a genotypic effect of >30% in cartilage, synovium and blood but only 3.9% in fat pad. These large differences imply intra-tissue differences in the nature of the mQTLs.

**Figure 3.**
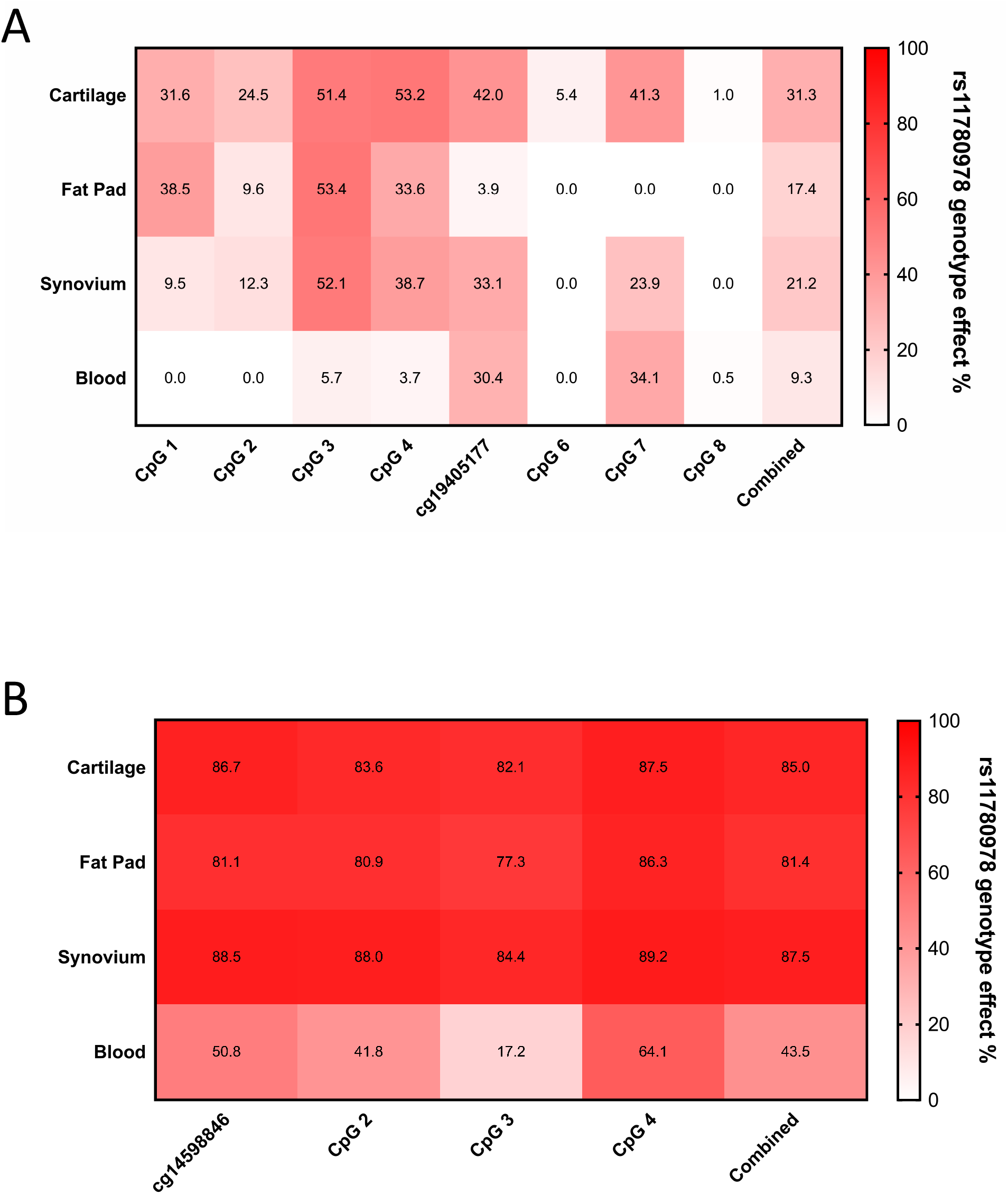
Heat maps displaying the percentage effect of genotype at rs11780978 upon methylation at CpG clusters cg19405177 (**A**) and cg14598846 (**B**) for the four patient tissues. Data analysis was performed using SAS JMP Statistical Data Visualization Software and heat maps produced in PRISM. Patients contributing data to the cg19405177 cluster 1: cartilage, n=81; fat pad, n=39; synovium, n=64; blood, n=43. Patients contributing data to the cg14598846 cluster: cartilage, n=103; fat pad, n=68; synovium, n=79; blood, n=54.

Overall, this analysis highlighted that the previously reported *PLEC* mQTLs are active in non-cartilaginous tissues and that the effects are stronger in joint tissues than blood. Although the genotypic effect of rs11780978 is greater in cartilage for the cg19405177 CpGs, this is not the case for the cg14598846 CpGs.

### The *PLEC* rs11780978 eQTL is active in synovium

Before testing for the *PLEC* rs11780978 eQTL by AEI analysis, we measured the relative expression of the gene in fat pad, synovium and blood along with cartilage as a comparator (Supplementary Figure 7). The qPCR data revealed that *PLEC* expression was equivalent between cartilage and synovium, was higher in fat pad and lower in blood.

We performed AEI as described previously^16^. In summary, we identified rs11780978 heterozygotes who were also heterozygous for the *PLEC* transcript SNP rs11783799 (r^2^=0.93; the OA risk conferring A allele of rs11780978 correlates with the G allele of rs11783799). We then used rs11783799 to quantify the relative allelic expression of *PLEC* in the compound heterozygotes. The blood cDNA samples consistently failed quality control, with the PCR replicates differing by >5% in all samples. This is likely a reflection of the lower expression of *PLEC* in this tissue and we could not therefore generate blood AEI data. We were able to generate AEI data for 19 fat pad and 20 synovium compound heterozygotes (Figure 4). AEI was detectable in both tissues and as for our previous study in cartilage^16^, the AEI ratios varied between patients, implying that other genetic polymorphisms can also modulate *PLEC* expression. Only in synovium did the direction of AEI significantly correlate with an allele of rs11783799, with reduced expression of *PLEC* correlating with the G allele of the SNP in the combined analysis (*P*=0.0054; Figure 4B). In fat pad, AEI was observed in a majority of the patients, but this did not correlate with a particular allele (*P*=0.31; Figure 4A).

**Figure 4.**
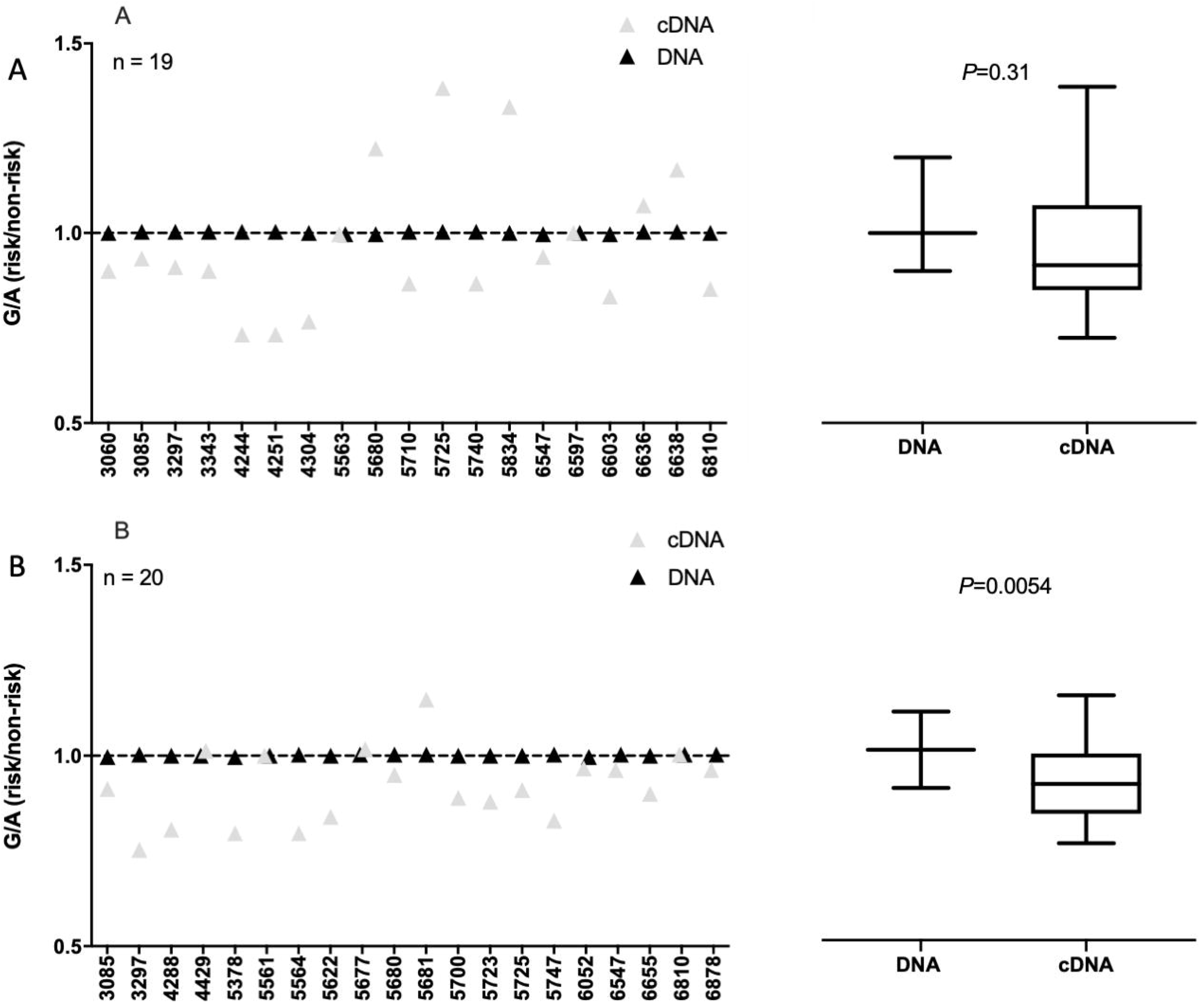
*PLEC* allelic expression imbalance (AEI) in fat-pad (**A**) and synovium (**B**). AEI analysis was conducted using transcript SNP rs11783799. The risk/non-risk allelic ratio is plotted, with a ratio <1 indicating decreased *PLEC* expression from the risk allele. For each patient, the mean of the DNA ratio (black, n=3 technical repeats) and the mean of the cDNA ratio (grey, n=3 technical repeats) is plotted. Numbers on the x-axis refer to the anonymised identification number assigned to patients at recruitment. The values for DNA and cDNA for all patients combined are represented as box plots to the right, in which the lines within the box represent the median, the box represents the 25th to 75th percentile, and the whiskers represent the maximum and minimum values. *P* values were calculated using Wilcoxon’s matched pairs signed rank test.

To emphasize the variability of AEI in tissues from the same patient, we next plotted the data for those patients in whom we had studied fat pad and synovium (Supplementary Figure 8). There were six such patients with clear variability in the G/A ratio between the two tissues tested for most of these, with patients 5680 and 5725 demonstrating particularly clear inter-tissue variability in the ratio; in both, the G allele is more highly expressed in fat pad but more lowly expressed in synovium relative to the A allele.

The significant AEI in synovium was in the same direction that we had observed in cartilage^16^, with the G allele of rs11783799, which marks the OA risk conferring A allele of rs11780978, demonstrating reduced *PLEC* expression. In that previous report, we had studied 19 cartilage compound heterozygotes and the mean G/A ratio was 0.89. The mean synovium G/A ratio we detected here was 0.92.

We next plotted the *PLEC* AEI data for the fat pad and synovium compound heterozygotes against their individual methylation data. In fat pad, there was no evidence of a significant correlation between methylation and AEI at cg19405177, cg14598846 or the additional 10 CpGs measured (all *P* values >0.05; Supplementary Figure 9). However, in synovium, three of the CpGs correlated with AEI; CpG3 from the cg19405177 cluster (*P*=0.049), and cg14598846 (*P*=0.039) and CpG2 (*P*=0.047) from the cg14598846 cluster (Supplementary Figure 10). These three CpGs had also shown meQTL activity in our previous cartilage study^16^. For CpG3 from the cg19405177 cluster, the same direction of effect is seen in cartilage and synovium, with increased methylation correlating with decreased AEI (the log_2_ AEI ratio moves towards zero with increasing methylation). However, for cg14598846 and CpG2 from the cg14598846 cluster, the effects are in opposite directions between cartilage and synovium; in cartilage, increased methylation correlated with decreased AEI whereas in synovium, increased methylation correlated with increased AEI (the log_2_ AEI ratio moves away from zero with increasing methylation).

In summary, the *PLEC* rs11780978 eQTL previously observed in cartilage also operates in synovium. Furthermore, we observed correlations between *PLEC* expression and CpG methylation in synovium. These meQTLs are shared with cartilage but are not necessarily operating in the same manner.

### Knock-down of *PLEC* and plectin elicits a response

Our previous data and the results generated here highlight that the OA risk conferring A allele of rs11780978 correlates with decreased expression of *PLEC* in cartilage and synovium. We modelled this effect by knocking-down plectin in an immortalised human MSC cell line. We chose to use this line because: 1) being an MSC line, it is biologically relevant to joint cells; 2) it expressed *PLEC* at a high level and; 3) it is readily transfectable and modifiable via CRISPR/Cas9^31^. We transduced it to constitutively express Cas9 and we then generated a 26bp frame-shift deletion of *PLEC* exon 3, with four replicates for deletion and four control replicates. The cells were cultured in monolayer for up to 14 days and the deletion was confirmed by PCR and Sanger sequencing, whilst a near-total plectin knock-out was observed by immunoblotting using an antibody that binds to the C-terminus of the protein (Figure 5A-C). The isolated RNA was then subjected to next-generation mRNA sequencing. This revealed alternate splicing occurring in *PLEC* in the deletion samples, with exon 3 being completely skipped in a subset of transcripts (Figure 5D). No exon skipping was seen in the control samples. The level of *PLEC* mRNA was 26% lower in the deletion samples compared to the controls (log_2_ fold change of −0.44; FDR *P*=0.001; Supplementary Table 2). In summary, this data implies that the Cas9-mediated deletion impacted on the transcription of *PLEC* and on the translation of *PLEC* mRNA into plectin.

**Figure 5.**
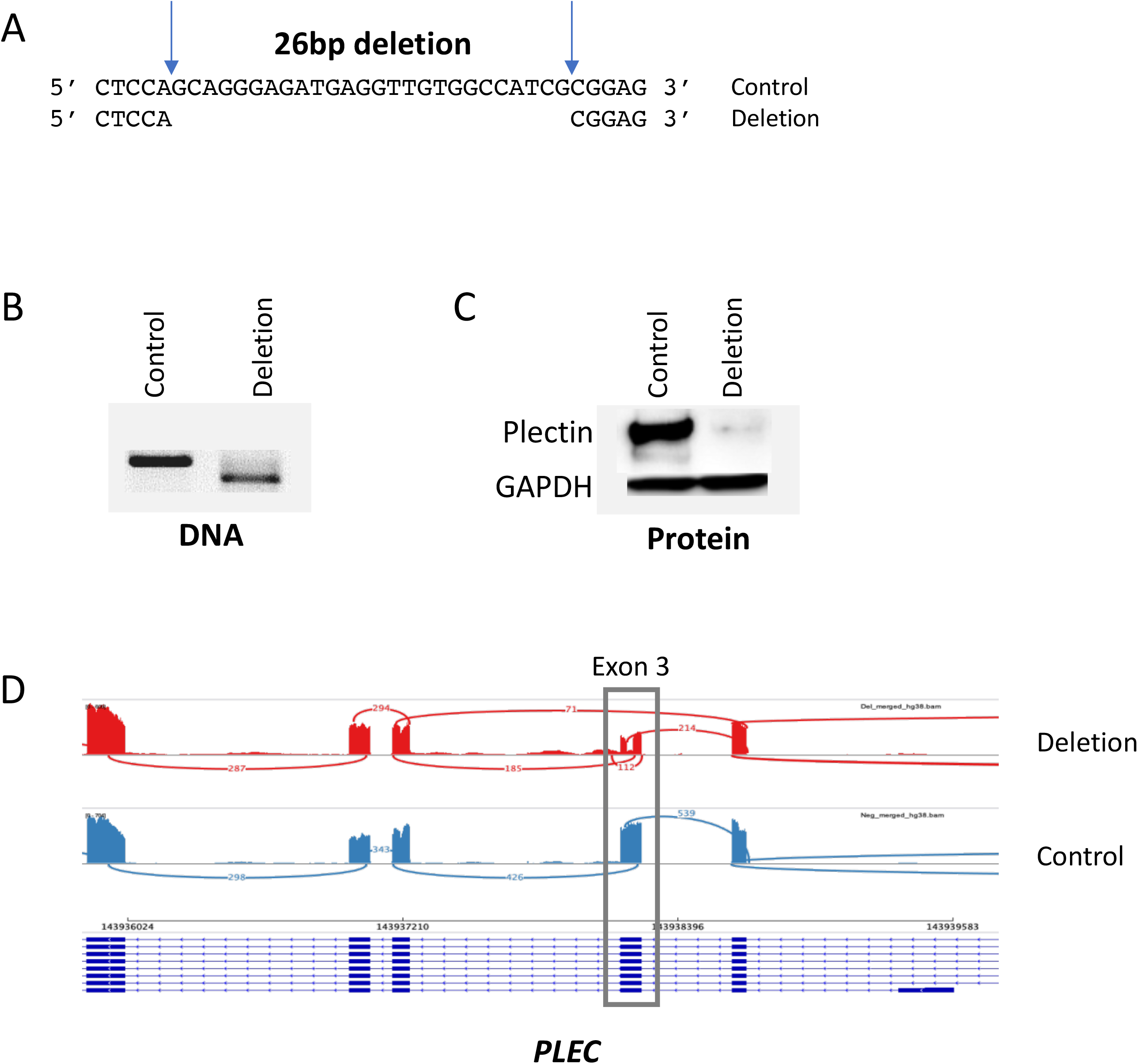
Knock-down of plectin using CRISPR/Cas9. (**A**) The 26bp sequence of *PLEC* exon 3 deleted. (**B**) Representative agarose gel image showing the PCR product for control (non-deleted) and deletion. (**C**) Representative immunoblot comparing plectin levels in control versus deletion, with GAPDH as a loading control. (**D**) Sashimi plot showing splicing of merged *PLEC* deletion (red) and control (blue) samples. Data were viewed on IGV genome browser. The multiple isoforms of *PLEC* are shown on the bottom of the panel. Read coverages show the presence of reads within exons. The lines illustrate the splicing between exons. The values on the lines show the number of junction spanning reads. The merged deletion track shows the absence of the 26bp region deleted within *PLEC* exon 3 (indicated by the box).

We next assessed the effect of this plectin knock-down on the transcriptome. In addition to *PLEC*, the deletion had a significant effect (FDR *P*<0.05) on the expression of 269 genes (Figure 6A and B, Supplementary Table 2). Pathway analysis revealed upregulation of processes involving immune regulation and protein post-translation modification (Figure 6C), and downregulation of Wnt signalling, metabolic pathways and glycosaminoglycan biosynthesis, amongst others (Figure 6D). In conclusion, this model system highlighted that depletion of *PLEC* and plectin impacted upon a range of pathways reported to play a critical role in cartilage biology and OA.

**Figure 6.**
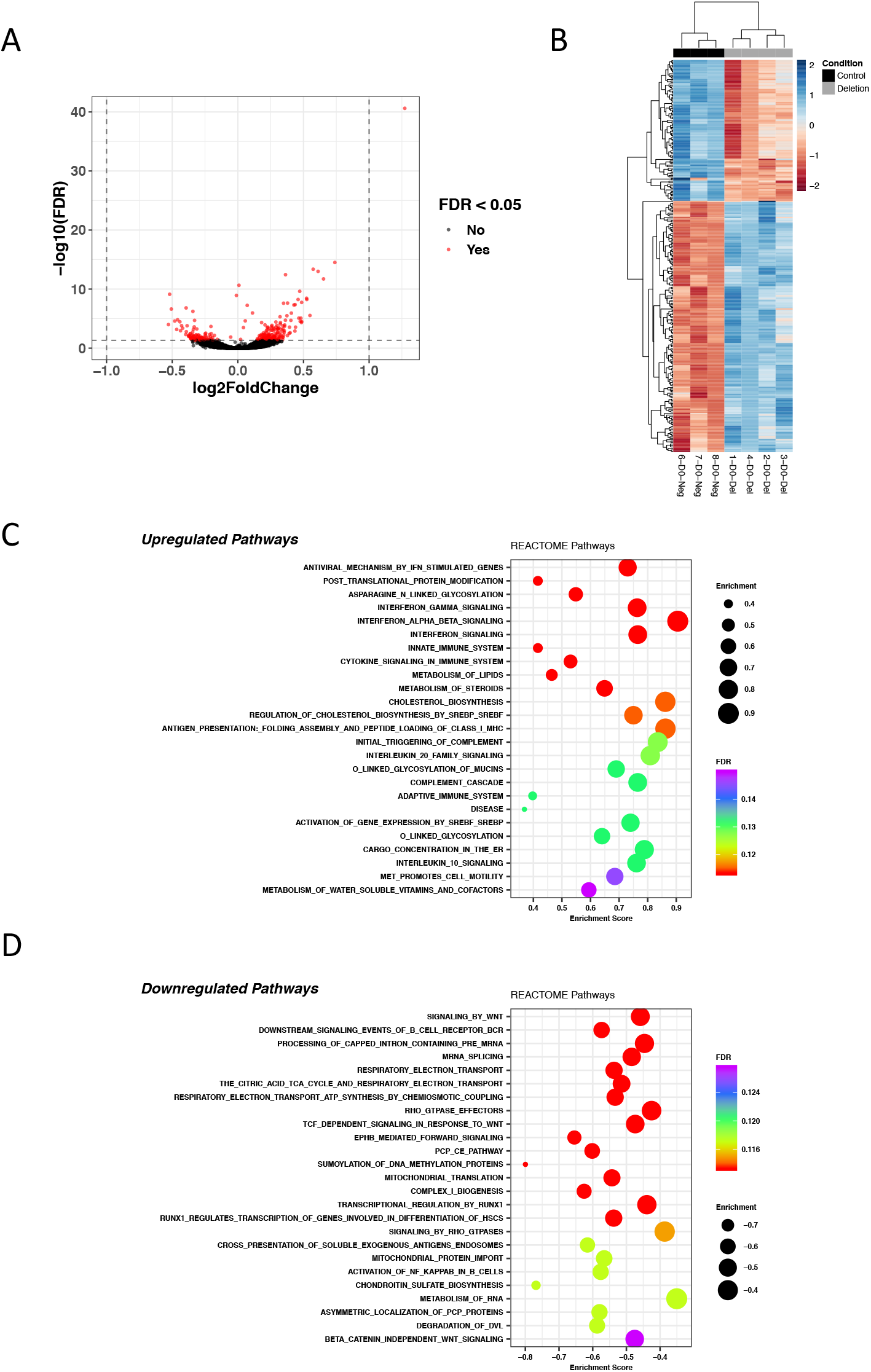
Differentially expressed genes and pathways. (**A**) Volcano plot illustrating significantly differentially expressed genes between *PLEC* deletion and control samples. The x-axis shows the log_2_ fold change between condition and the y-axis shows the −log_10_(FDR). Genes with an FDR *P*<0.05 are shown in red. (**B**) Heatmap of significantly differentially expressed genes between *PLEC* deletion and control. Normalized counts were used for plotting. Hierarchical clustering was used to cluster samples and genes. The heatmap shows that samples cluster by condition (D0-Del, deletions; D0-Neg, controls). (**C**) Top 25 significantly upregulated Reactome pathways found using GSEA. (**D**) Top 25 significantly downregulated Reactome pathways found using GSEA. All genes were ranked using signed log_10_(*P*-value) and this pre-ranked list used as input into GSEA to query enrichment of Reactome pathways.

## Discussion

The mapping of risk loci for polygenic traits is now a relatively straightforward procedure, with the next major step in complex trait analysis being the transition from association signal to functional characterization^32^. For OA, considerable strides have been made in this regard in recent years^33–35^. Our interest is in the epigenetic dimension of OA genetic risk and especially the role of DNA methylation. This is the best characterized and most stable epigenetic modification and modulates gene expression by disrupting the binding of transcription factors and by facilitating the binding of methyl-binding proteins, which initiate compaction of chromatin and gene silencing^36^. Last year, we demonstrated that the OA association signal marked by SNP rs11780978 correlated with methylation of CpGs within *PLEC* and with variable expression of the gene^16^. These analyses were chondro-centric and here, we set out to determine whether these functional effects were more widespread. If so an exclusive focus on cartilage as a target tissue of the rs11780978 association signal would not be merited.

We discovered that the rs11780978-*PLEC* eQTLs, mQTLs and meQTLs that are active in cartilage are also active in synovium, but with some differences; the genotypic effect is substantially greater in cartilage for the cg19405177 cluster of CpGs, whilst the cg14598846 meQTLs act in opposite directions between the two tissues. Further investigation in additional cohorts should confirm whether both the commonalty and discord of the effects that we have observed between cartilage and synovium are robust. In fat pad, we observed the mQTLs but *PLEC* AEI did not correlate with rs11780978 genotype. Instead, the OA risk-conferring A allele of rs11780978 demonstrated increased and decreased expression relative to the non-risk allele across the 19 fat pad samples studied, with no overall significant difference when the data was combined. There are therefore eQTLs operating on *PLEC* in fat pad, but these are unrelated to rs11780978. These distinctions point toward regulatory variability in gene expression mediated by the same association between joint tissues. In blood, the mQTLs were detectable but their genotypic effect was much weaker in comparison to the three joint tissues. Such variable strengths of mQTLs effects between cells from different lineages are expected^37^. We were unable to test blood for *PLEC* eQTLs due to the low expression level of the gene. In summary, we observed the *PLEC* mQTLs across all four tissues tested with the joint tissues more similar to each other than to blood. This was exemplified by the mQTL genotypic effects being much stronger in joint tissues. Since synovium was subject to eQTLs and meQTLs, it too should be investigated as a target of the rs11780978 association signal.

Finally, we modelled the effect of the rs11780978 risk-conferring allele by knocking down plectin in a cell line. *PLEC* mRNA was detectable in the RNA-sequencing data but at a significantly lower level compared to control. The knock-down was very efficient at the protein level, with a near total absence of plectin observed. The knock-down impacted on a range of cellular pathways. Of particular note, both the innate and the acquired immune response showed an upregulation, an effect that has been reported on previously in OA for both cartilage and synovium^38–39^. Downregulation of Wnt signalling was also of interest, as this pathway is critical to cartilage homeostasis^33,40^. Furthermore, there is growing evidence of an interplay between Wnt signalling and inflammation in joint tissues^41^. Our model system did not include mechanical load, a force which plectin allows cells to respond to^18–20^. Aberrant mechanical load, including only moderate change, can elicit an inflammatory response that has been termed “mechanoflammation”^42^. Our data suggests that even in the absence of load, the loss of plectin is eliciting this response. A future model incorporating load may therefore reveal yet more insight into a potential key role of plectin as a regulator of inflammation in response to mechanical load and joint tissue injury in OA. This model should incorporate plectin deficient differentiated joint cells, rather than the undifferentiated MSCs investigated here, to more accurately recapitulate the articulating joint. Furthermore, and despite our inability to undertake *PLEC* AEI on blood cells, we should not exclude circulating immune cells as potential targets of the rs11780978 association signal, particularly in light of the altered expression of plectin in macrophages in response to immunomodulatory agents^43^.

In conclusion, our investigations have highlighted the interplay between OA genetic risk, DNA methylation and gene expression, and reveal clear differences in these effects for tissues from the same diseased joint. Furthermore, by recapitulating the effect of the risk locus at *PLEC*, we have linked OA genetic susceptibility with a number of cellular pathways, thus providing mechanistic insight into OA aetiology.

## Author Contributions

AKS and JL designed the study. AKS, IMJH, MT and SJR generated the experimental data. AKS, IMJH, KC and EP analysed the data and prepared the manuscript figures. AKS, IMH, and JL prepared the main manuscript text. DJD provided patient samples for analyses. All authors were involved in drafting the article or revising it critically for intellectual content, and all authors approved the final version to be published.

## Acknowledgements

Dr Colin Shepherd provided input into data interpretation and reviewed the manuscript.

## Data Availability

All data are available from the corresponding authors upon request. Raw RNA-seq data has been deposited into the NCBI GEO datasets repository with accession number GSE143725.

---

**Supplementary Figure 1.** The location in *PLEC* of rs11780978 and the analysed CpGs. The *PLEC* gene is shown in blue, exons are boxes/vertical lines, and the direction of transcription is from right to left. The cg19405177 assay captures a total of 8 CpGs, the cg14598846 assay a total of four. cg19405177 and cg14598846 are marked as red circles, the additional CpGs as black circles. The physical position of the CpGs on chromosome 8 (hg19 assembly, UCSC Genome Browser) is shown.

**Supplementary Figure 2.** Association between rs11780978 genotype and methylation at CpG clusters cg19405177 (**A**) and cg14598846 (**B**) in cartilage DNA. *P* values were calculated using the Kruskal-Wallis test. Horizontal lines and error bars show the mean ± SEM. n = the number of patients providing data per CpG site.

**Supplementary Figure 3.** Association between rs11780978 genotype and methylation at CpG clusters cg19405177 (**A**) and cg14598846 (**B**) in fat pad DNA. *P* values were calculated using the Kruskal-Wallis test. Horizontal lines and error bars show the mean ± SEM. n = the number of patients providing data per CpG site.

**Supplementary Figure 4.** Association between rs11780978 genotype and methylation at CpG clusters cg19405177 (**A**) and cg14598846 (**B**) in synovium DNA. *P* values were calculated using the Kruskal-Wallis test. Horizontal lines and error bars show the mean ± SEM. n = the number of patients providing data per CpG site.

**Supplementary Figure 5.** Association between rs11780978 genotype and methylation at CpG clusters cg19405177 (**A**) and cg14598846 (**B**) in blood DNA. *P* values were calculated using the Kruskal-Wallis test. Horizontal lines and error bars show the mean ± SEM. n = the number of patients providing data per CpG site.

**Supplementary Figure 6.** Methylation at CpG clusters cg19405177 (**A**) and cg14598846 (**B**) for the four patient tissues without stratification by rs11780978 genotype. *P* values were calculated using Kruskal-Wallis with Dunn’s multiple comparisons. Horizontal lines and error bars show the mean ± SEM. n = the number of patients providing data per tissue and at each CpG site. \**P*◻≤◻0.05; \*\**P*◻≤◻0.01; \*\*\**P*◻≤◻0.001 \*\*\*\**P*◻≤◻0.0001.

**Supplementary Figure 7.** Expression of *PLEC* in tissue samples from OA patients. *PLEC* mRNA levels were measured by qPCR in cartilage (n=10), fat pad (n=9), synovium (n=10) and blood (n=10). Horizontal lines and error bars show the mean ± SEM. *P* values were calculated using a Mann-Whitney 2-tailed exact test. \*\**P*◻0.01; \*\*\**P*◻≤◻0.001

**Supplementary Figure 8.** The six patients with AEI data for fat pad (FP) and synovium (SYN). The risk/non-risk allelic ratio is plotted, with a ratio <1 indicating decreased *PLEC* expression from the risk allele. For each patient, the mean of the DNA ratio (black, n=3 technical repeats) and the mean of the cDNA ratio (grey, n=3 technical repeats) is plotted. Numbers on the x-axis refer to the anonymised identification number assigned to patients at recruitment.

**Supplementary Figure 9.** Methylation expression quantitative trait locus analyses in fat pad. *PLEC* log_2_ allelic expression imbalance (AEI) ratios were plotted against methylation at all CpGs in the cg19405177 (**A**) and cg14598846 (**B**) clusters. The square of the correlation coefficient (r^2^) and *P* values were calculated by linear regression analysis using a standard least squares model.

**Supplementary Figure 10.** Methylation expression quantitative trait locus analyses in synovium. *PLEC* log_2_ allelic expression imbalance (AEI) ratios were plotted against methylation at all CpGs in the cg19405177 (**A**) and cg14598846 (**B**) clusters. The square of the correlation coefficient (r^2^) and *P* values were calculated by linear regression analysis using a standard least squares model.

