## Supplementary figures and images for "Multi-tissue epigenetic analysis of the osteoarthritis susceptibility locus mapping to the plectin gene *PLEC*"

### Supplementary Figure 1

#### Supplementary Figure 1

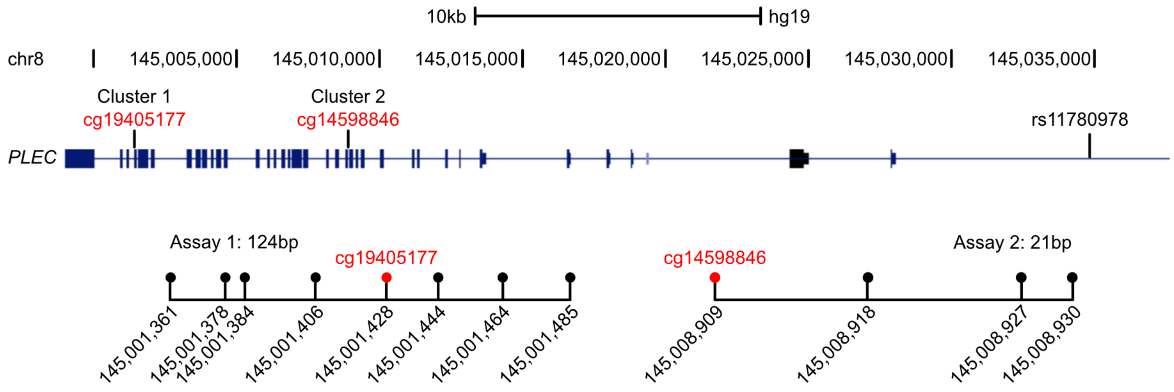

### Supplementary Figure 2

#### Supplementary Figure 2

A

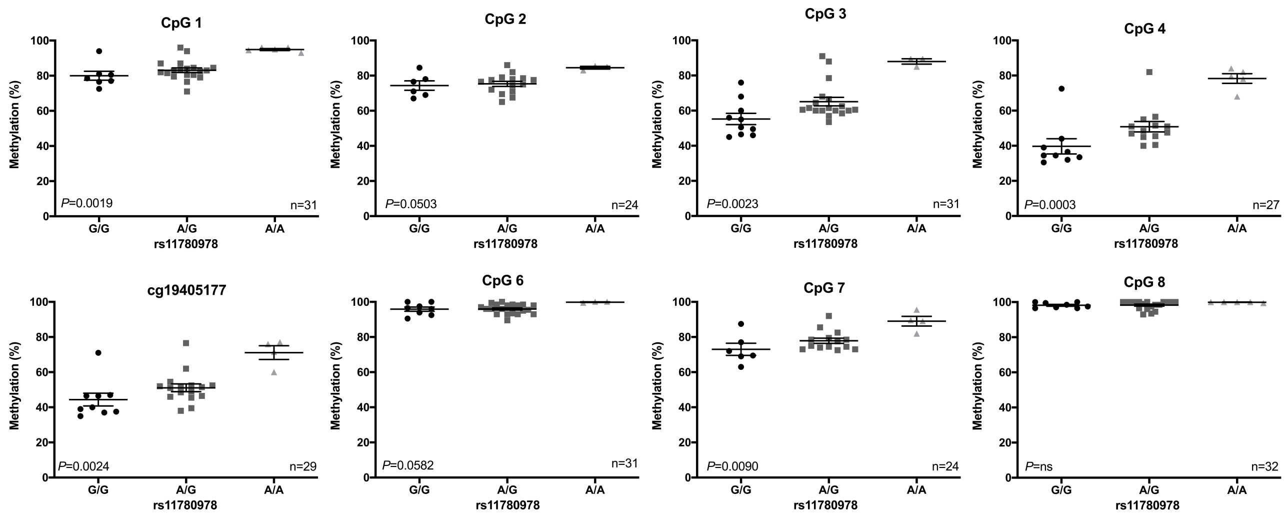


B

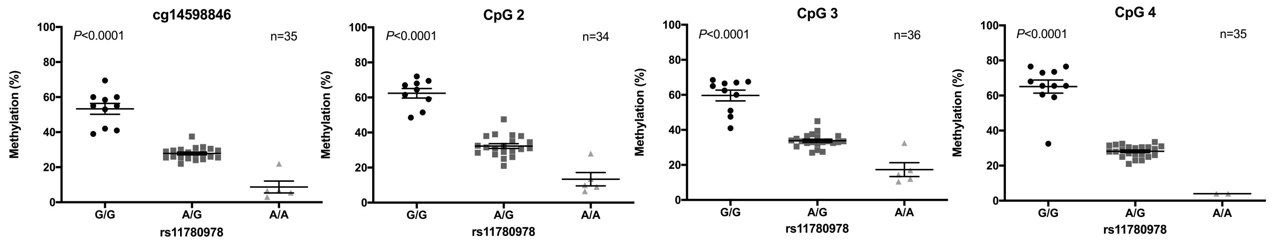

### Supplementary Figure 3

##### Supplementary Figure 3

A

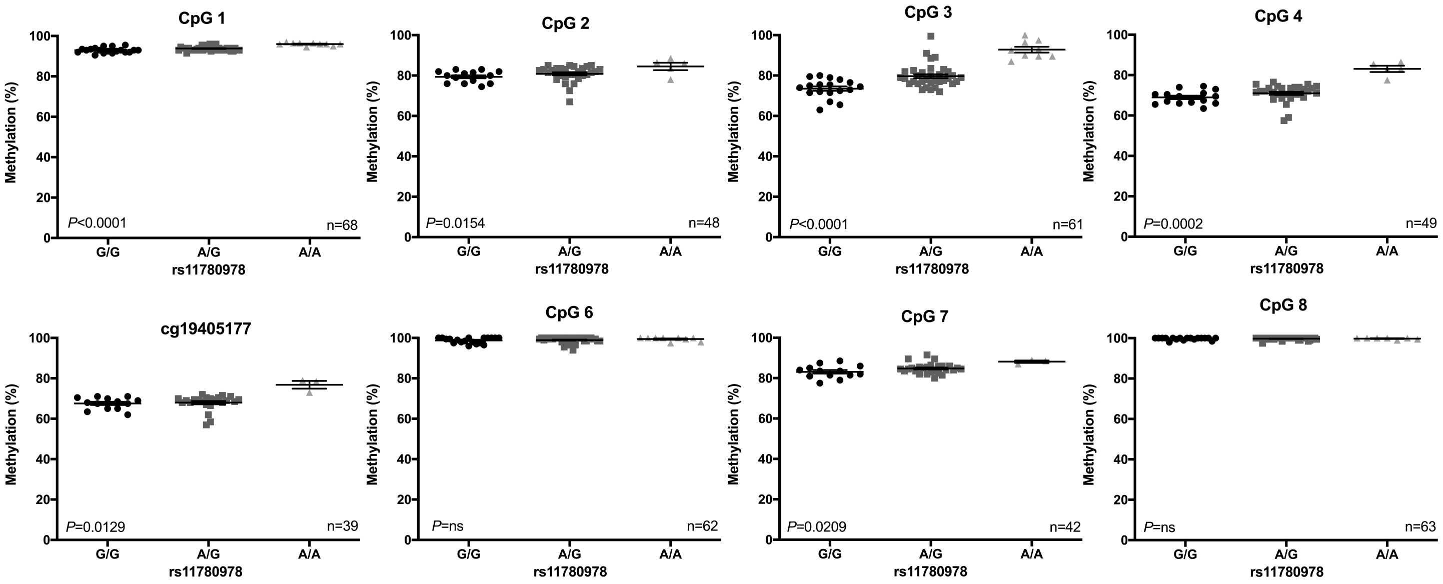


B

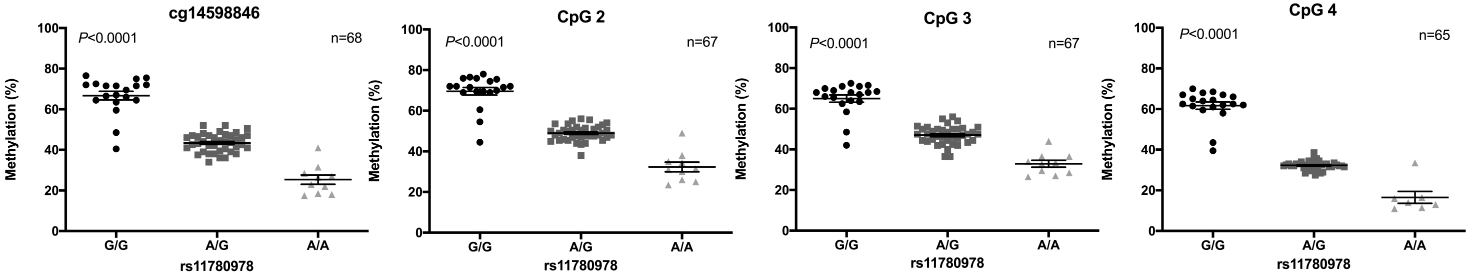

### Supplementary Figure 4

##### Supplementary Figure 4

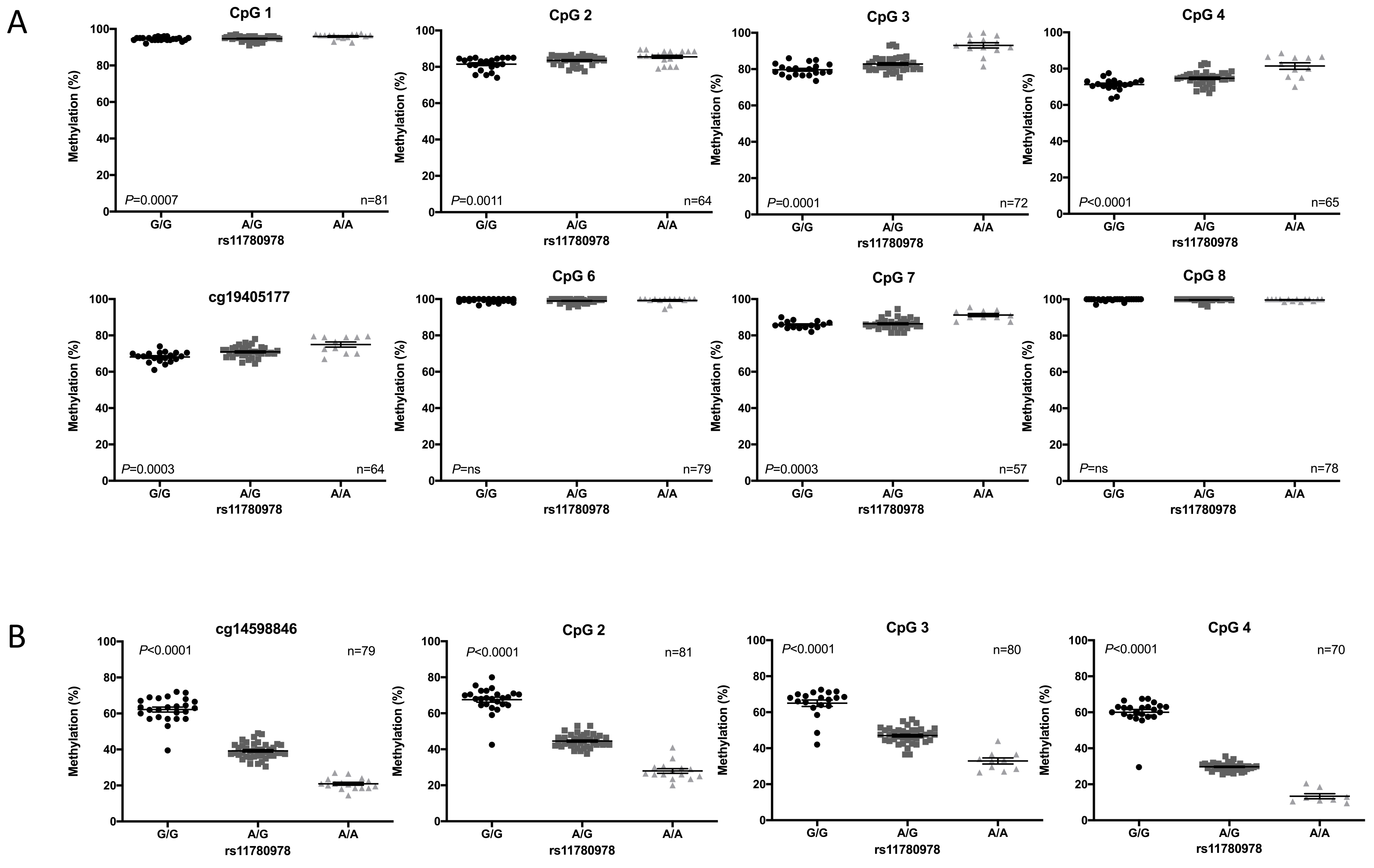

### Supplementary Figure 5

##### Supplementary Figure 5

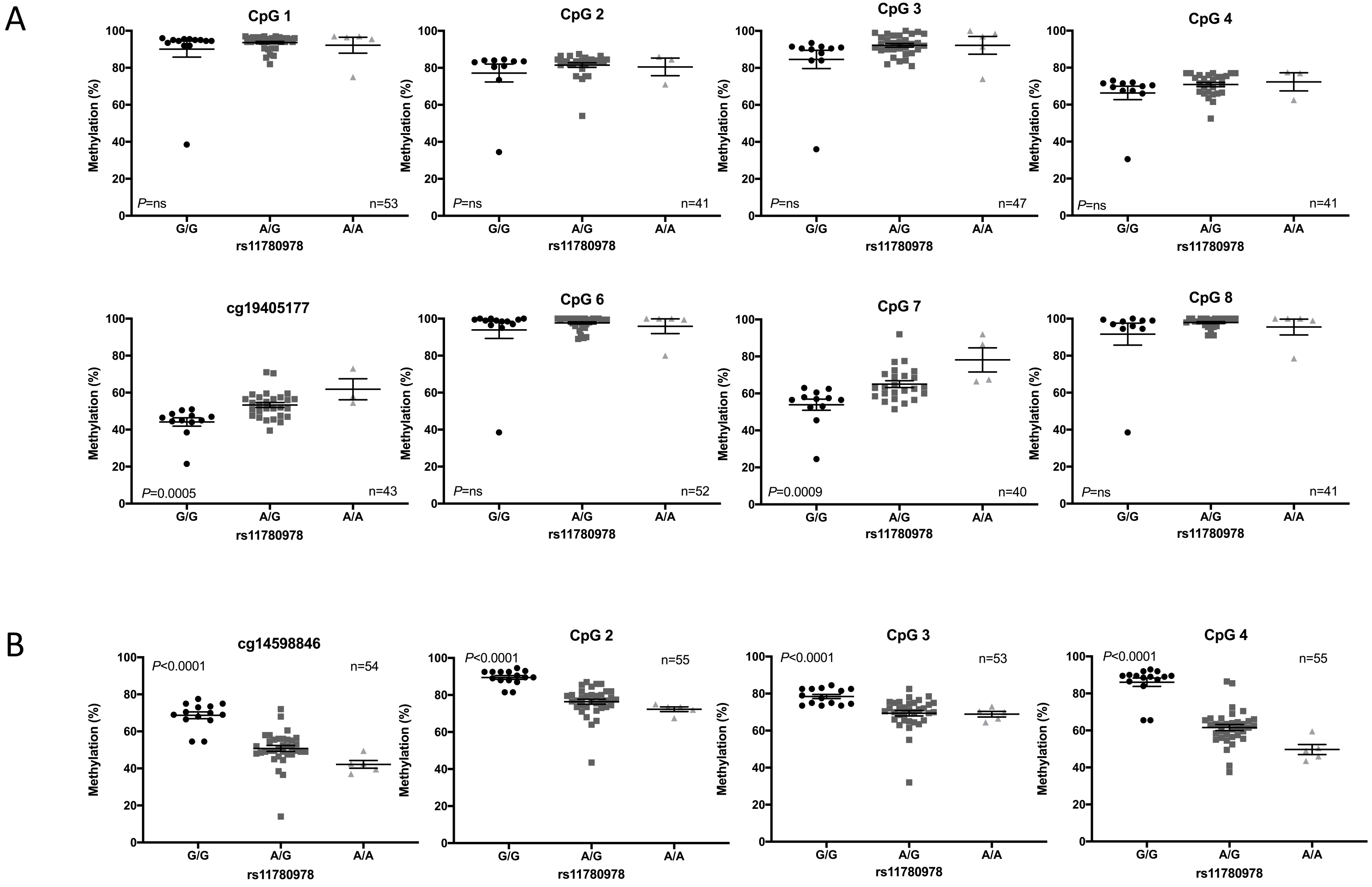

### Supplementary Figure 6

##### Supplementary Figure 6

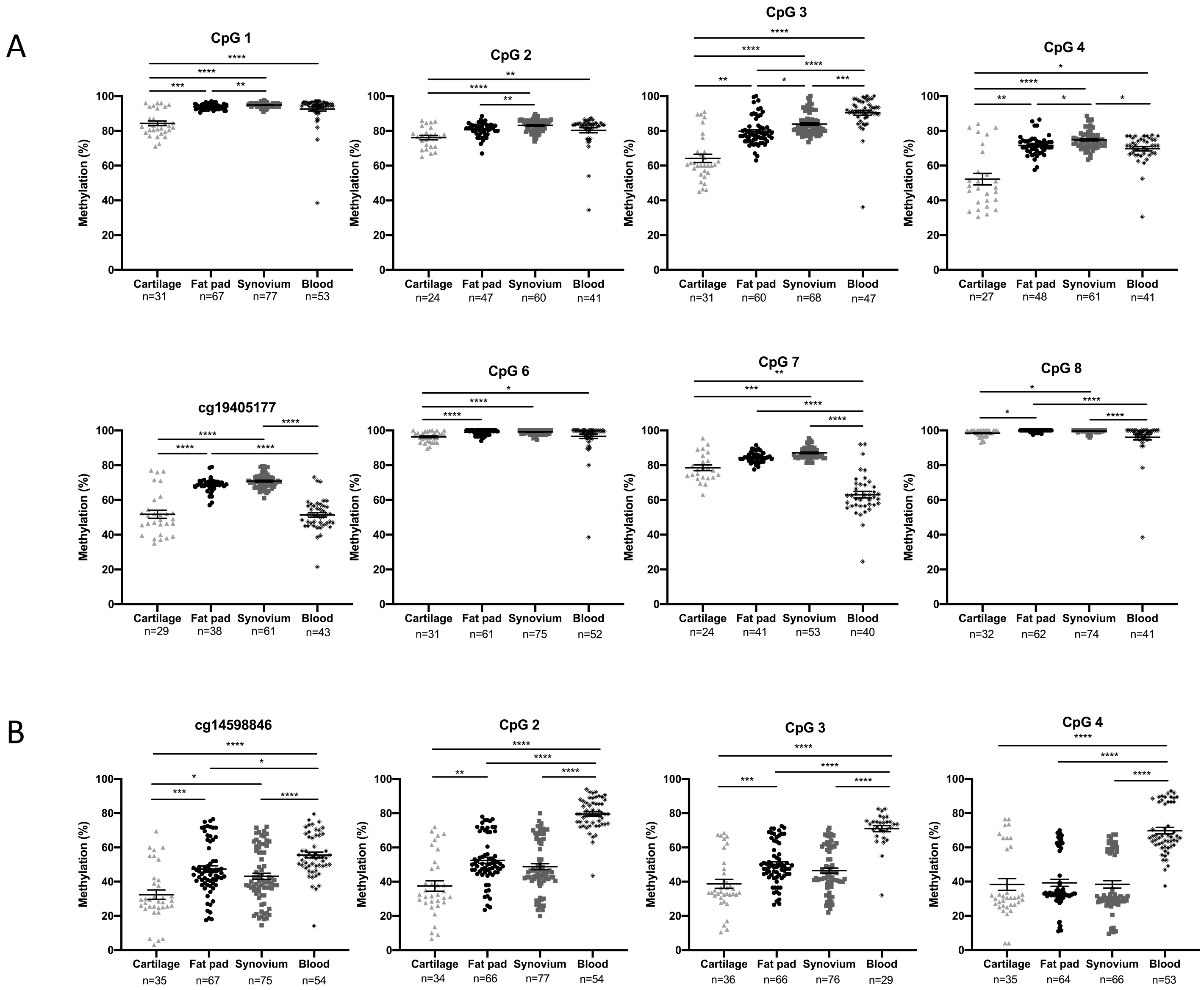

### Supplementary Figure 7

**Supplementary Figure 7**

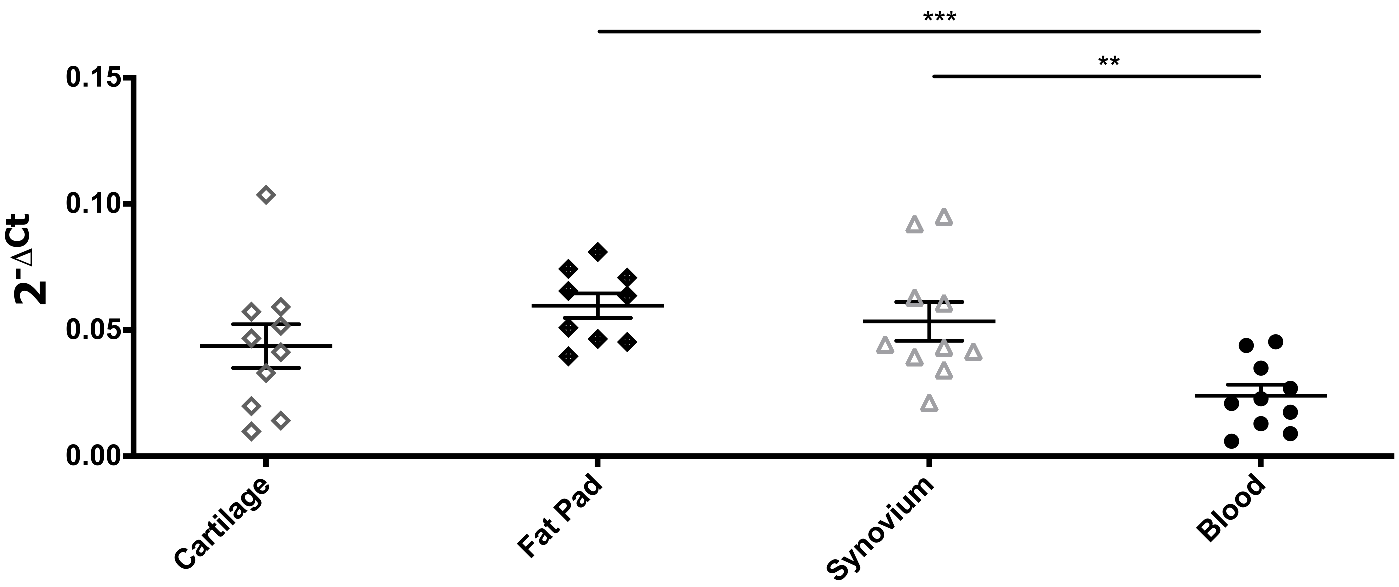

### Supplementary Figure 8

**Supplementary Figure 8**

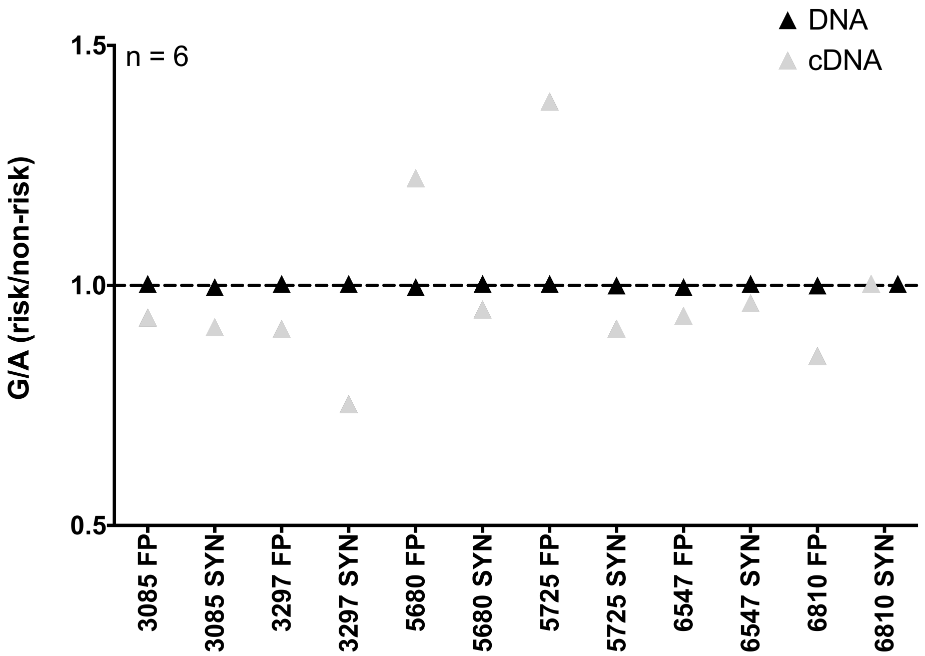

### Supplementary Figure 9

##### Supplementary Figure 9

A

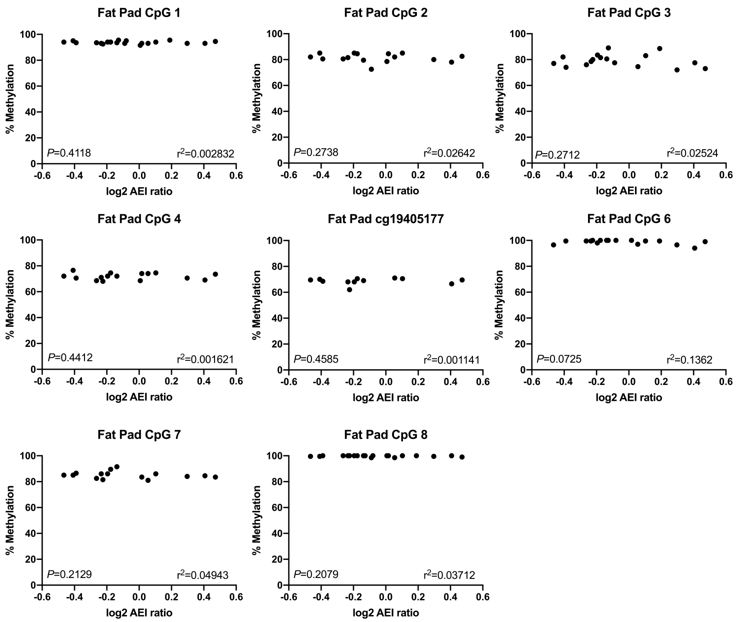


B

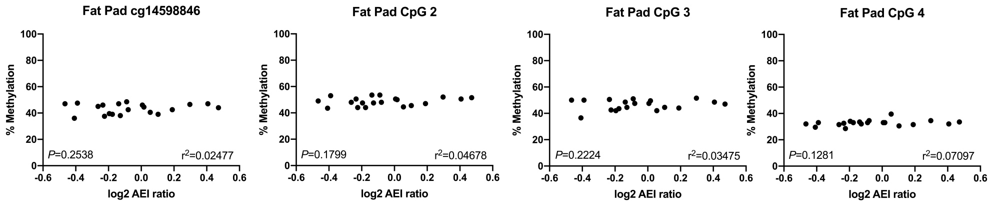

### Supplementary Figure 10

##### Supplementary Figure 10

A

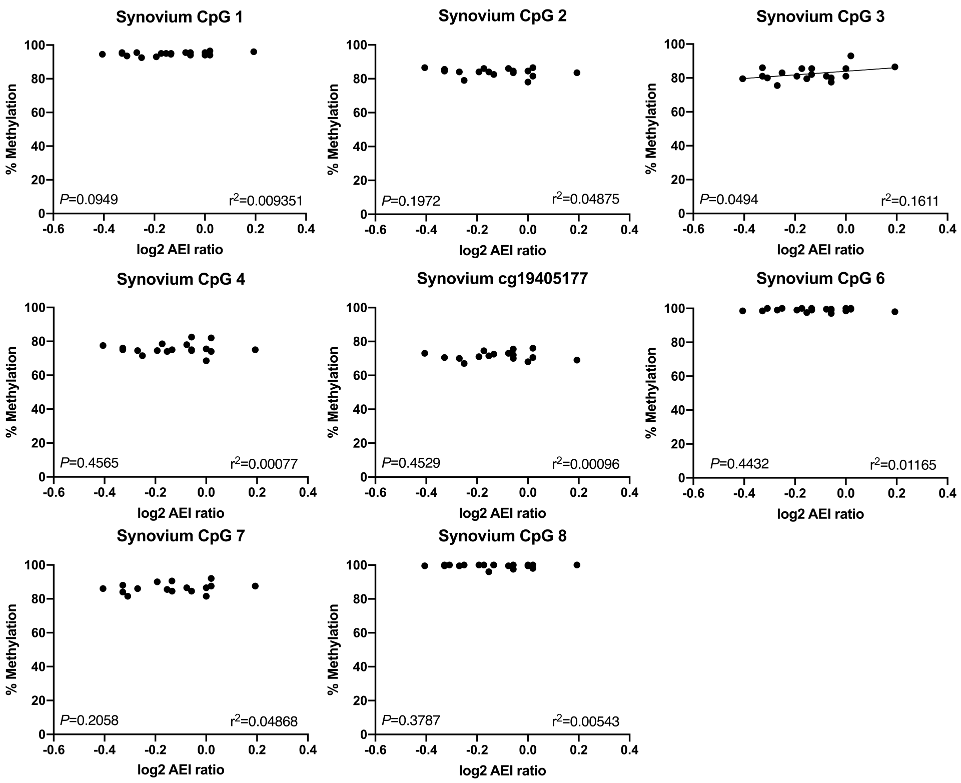


B

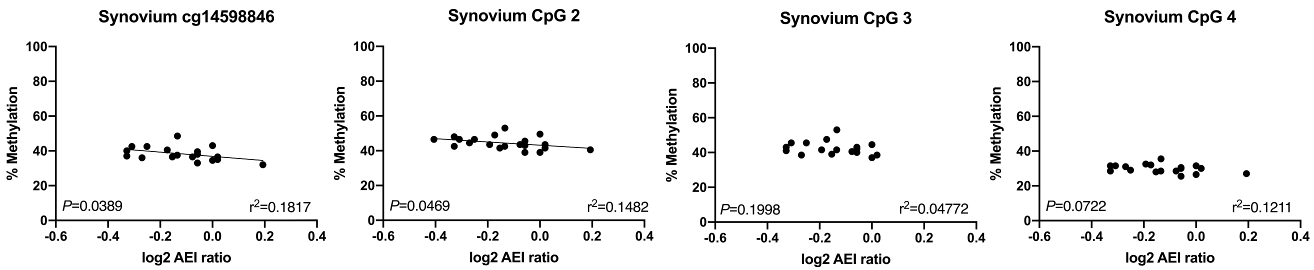
